# Inhibition of Angiotensin II Dependent AT1a Receptor Stimulation Attenuates Thoracic Aortic Pathology in Fibrillin-1^C1041G/+^ Mice

**DOI:** 10.1101/2020.06.01.127670

**Authors:** Jeff Z. Chen, Hisashi Sawada, Jessica J. Moorleghen, Michael K. Franklin, Deborah A. Howatt, Mary B. Sheppard, Adam E. Mullick, Hong S. Lu, Alan Daugherty

## Abstract

**Graphic Abstract:** 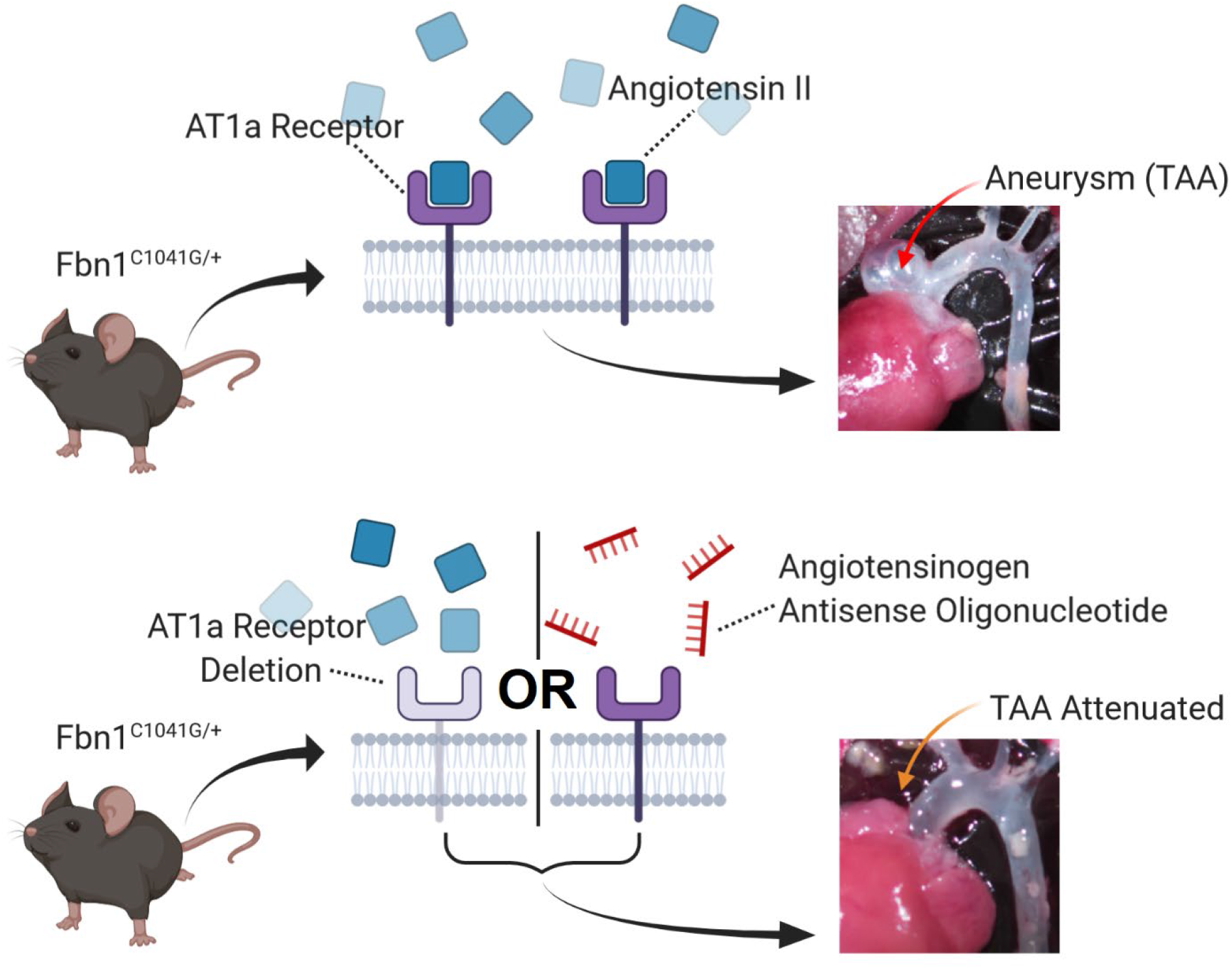

**Objective:** A cardinal feature of Marfan syndrome is thoracic aortic aneurysm (TAA). The contribution of ligand-dependent stimulation of angiotensin II receptor type 1a (AT1aR) to TAA progression remains controversial because the beneficial effects of angiotensin receptor blockers have been ascribed to off-target effects. This study used genetic and pharmacologic modes of attenuating angiotensin receptor and ligand, respectively, to determine their roles on TAA in mice with fibrillin-1 haploinsufficiency (Fbn1^C1041G/+^).

**Approach and Results:** TAA in Fbn1^C1041G/+^ mice were determined in both sexes and found to be strikingly sexual dimorphic. Males displayed progressive dilation over 12 months while ascending aortic dilation in Fbn1^C1041G/+^ females did not differ significantly from wild type mice. To determine the role of AT1aR, Fbn1^C1041G/+^ mice that were either +/+ or −/− for AT1aR were generated. AT1aR deletion reduced progressive expansion of ascending aorta and aortic root diameter from 1 to 12 months of age in males. Medial thickening and elastin fragmentation were attenuated. An antisense oligonucleotide against angiotensinogen (AGT-ASO) was administered to male Fbn1^C1041G/+^ mice to determine the effects of angiotensin II depletion. AGT-ASO administration, at doses that markedly reduced plasma AGT concentrations, attenuated progressive dilation of the ascending aorta and aortic root. AGT-ASO also reduced medial thickening and elastin fragmentation.

**Conclusions:** Genetic approaches to delete AT1aR and deplete AngII production exerted similar effects in attenuating pathology in the proximal thoracic aorta of male Fbn1^C1041G/+^ mice. These data are consistent with ligand (AngII) dependent stimulation of AT1aR being responsible for aortic disease progression.

**Highlights:**

- Profound sexual dimorphism of aortic disease occurs in Fbn1^C1041G/+^ mice, with female mice being more resistant and male mice being more susceptible.
- Inhibition of the AngII-AT1aR axis attenuates aortic pathology in male Fbn1^C1041G/+^ mice.
- Antisense oligonucleotides targeting angiotensinogen deplete plasma angiotensinogen and attenuate thoracic aortic aneurysms.

## Introduction

Marfan syndrome is an autosomal dominant genetic disorder associated with thoracic aortic aneurysm (TAA) that enhance the risk for aortic rupture due to loss of aortic integrity.^1^ The disease is caused by mutations in fibrillin-1; a protein incorporated into the microfibrils that decorate elastic fibers.^2^ To gain insight into the mechanisms of the disease, mice have been developed with a heterozygous expression of the C1041G mutation of the mouse fibrillin-1 protein, which is analogous to the C1039Y mutation in humans.^3^ These mice have a haploinsufficiency of fibrillin-1 and mimic some pathologies present in patients with Marfan syndrome including progressive expansion of the proximal thoracic aorta.

The renin angiotensin system has been invoked as a mediator of TAA in patients with Marfan syndrome.^4^ Experimental evidence for the role of the renin angiotensin system has been based predominantly on the observation that losartan inhibits aortic pathology in mice. This was demonstrated initially in Fbn1^C1041G/+^ mice administered losartan starting at the prenatal phase of life.^5^ Additionally, it has been consistently demonstrated that losartan reduces aortic expansion in many other mouse models of TAA.^6–11^ However, losartan has many well-characterized effects independent of AT1 receptor antagonism potentially could compromise interpretation as a pharmacological tool to specifically study AT1 receptors.^12^ Indeed, the benefit of losartan in inhibiting aortic root dilation in Fbn1^C1041G/+^ mice has been attributed to effects such as TGF-β antagonism or nitric oxide synthase stimulation.^5, 6^ To overcome the limitations of pharmacological approaches, there is a critical need to determine the role of AT1aR using genetic deletion to specifically ascribe a function of AT1 receptors in general and AT1aR specifically.

Additionally, the mode by which AT1 receptors become activated in Marfan syndrome is uncertain. While activation of AT1 receptors is commonly due to engagement of the ligand, angiotensin II (AngII), the pathway can also be activated by conformational changes of the receptor during cell stretch of myocytes and vascular smooth muscle cells.^13–15^ This stretch activation of AT1 receptor can be inhibited by specific pharmacological antagonists of the receptor. Based on studies in angiotensinogen deficient mice, dilated cardiomyopathy in the fibrillin-1 hypomorphic model of Marfan syndrome has been attributed to this AngII-independent activation of AT1aR.^16^ The relative role of receptor activation of ligand versus stretch has not been evaluated in vascular disease.

The aim of the present study was to define the contribution of ligand-dependent activation of AT1aR to the progressive expansion of the proximal thoracic aorta in Fbn1^C1041G/+^ mice. Aortic diameters were measured for a 1 year interval using a standardized ultrasound protocol.^17, 18^ In accord with current guidelines, the study was performed in both sexes of these mice. These studies demonstrated a strong sexual dimorphism with greater expansion in Fbn1^C1041G/+^ males and minimal progressive expansion in Fbn1^C1041G/+^ females. Surprisingly, this disparity has not been reported previously. The role of AT1aR was determined subsequently using male mice with global AT1aR deletion. The role of the ligand was determined using an angiotensinogen antisense oligonucleotide (AGT-ASO) that depleted the unique precursor of AngII. This study demonstrated the importance of ligand-dependent activation of AT1aR to progression of aortic pathology.

## Methods

Tabulated data supporting the findings of this study are available as a supplemental document. Raw data supporting the findings of this study are available from the corresponding author upon reasonable request.

### Mice

Studies were performed in accordance with recommendations for design and reporting of animal aortopathy studies.^19, 20^ Studies were performed using littermate controls. Mice and genealogy were tracked with Mosaic Vivarium Laboratory Animal Management Software (Virtual Chemistry). Male and female AT1aR deleted (AT1aR^−/−^) (stock #002682) and Fbn1^C1041G/+^ (stock #012885) mice were obtained from The Jackson Laboratory. Male AT1aR heterozygous (AT1aR^+/−^) x Fbn1^C1041G/+^ were bred with female AT1aR^+/−^ x fibrillin-1 wild type (Fbn1^+/+^) mice to generate four experimental groups per sex: male and female AT1aR wild type (AT1aR^+/+^) x Fbn1^+/+^, AT1aR^−/−^ x Fbn1^+/+^, AT1aR^+/+^ x Fbn1^C1041G/+^, and AT1aR^−/−^ x Fbn1^C1041G/+^ mice. Littermates were separated by sex and genotypes and were randomized when housing mice after weaning. For AGT ASO experiments, 2-month-old male Fbn1^C1041G/+^ mice were procured from The Jackson Laboratory and randomized into experimental groups using a random number generator. Mice were checked daily for health, and necropsy was performed to adjudicate cause of death. Mice were housed up to 5 per cage and maintained on a 14:10 hour light:dark cycle. Mice were fed Teklad Irradiated Global 18% Protein Rodent Diet # 2918 *ad libitum* and allowed *ad libitum* access to water via a Lixit system. Bedding was provided by P.J. Murphy (Coarse SaniChip) and changed weekly during the study. Cotton pads were provided as enrichment. The room temperature was maintained at 21°C and room humidity was maintained at 50%. All protocols were approved by University of Kentucky IACUC.

### Genotyping

Mice were genotyped twice using tail tissue. Group allocation was based on genotyping performed after weaning at postnatal day 28 and again after study termination to verify genotypes. AT1aR deletion was assayed using forward primer (5’-AAATGGCCCTTAACTCTTCTACTG-3’) and reverse primer (5’-ATTAGGAAAGGGAACAGGAAGC-3’) covering a neo cassette that disrupts AT1aR spanning bps 110-635. The neo cassette removed approximately 0.5 kb and inserted approximately 1 kb of neo gene. AT1aR^+/+^ generated a 631 bp product. AT1aR^−/−^ generated a ~1.1 kbp product. Fbn1^C1041G/+^ was assayed using forward primer (5’-CTCATCATTTTTGGCCAGTTG-3’) and reverse primer (5’-GCACTTGATGCACATTCACA-3’) covering a single loxP intronic sequence within intron 24 which is not present in wild type mice. The protocol used was as described by The Jackson Laboratory. Fbn1^+/+^ generates a 164 bp product. Fbn1^C1041G/+^ generates a 212 bp product. Post-termination validation genotyping was performed by Transnetyx.

### Antisense Oligonucleotides

Scrambled control ASO (5’-GGCTACTACGCCGTCA-3’) and AGT ASO (5’-ATCATTTATTCTCGGT-3’) were provided by Ionis Pharmaceuticals. ASOs were 3-10-3 2’ – 4’ constrained ethyl gapmers that have demonstrated improved potency and tolerability versus locked nucleic acid or 2’-O-methoxyethyl gapmers.^21^ Mice were randomized to study group using a random number generator. Two-month-old male Fbn1^C1041G/+^ mice were administered control ASO or AGT ASO (80 mg/kg) subcutaneously at day 1 and 3 of study. Mice were maintained on subcutaneous control ASO or AGT ASO (40 mg/kg) every 7 days for the remainder of the study.

### Ultrasound Measurements

Ultrasound was performed by standardized protocols that have been as described previously.^17, 22^ Briefly, mice were anesthetized using inhaled isoflurane (2-3% vol/vol) and maintained at a heart rate of 450-550 beats per minute during image capture to reduce anesthesia exposure and maintain consistent heart rate between animals (Somnosuite, Kent Scientific). The order by which mice were subject to ultrasound was randomized. Ultrasound images were captured using a Vevo 3100 system with a 40 MHz transducer (Visualsonics). Images captured were standardized according to two anatomical landmarks: the innominate artery branch point and aortic valves. The largest luminal ascending aortic diameter between the sinotubular junction and the innominate artery were measured in end-diastole over three cardiac cycles by two independent observers.

### Measurement of in situ Aortic Diameters

Mice were terminated by overdose of ketamine:xylazine followed by cardiac puncture and saline perfusion. The order in which mice were terminated was randomized. Aortas were dissected away from surrounding tissue and Optimal Cutting Temperature Compound (Sakura Finetek) was introduced into the left ventricle to maintain aortic patency. A black plastic sheet was inserted beneath the aorta and heart to increase contrast and facilitate visualization of aortic borders. Aortas were imaged using a Nikon SMZ800 stereoscope and measurements were recorded using NIS-Elements AR 4.51 software (Nikon Instruments Inc.). Ascending aortic diameters were measured at the largest width perpendicular to the vessel.

### Histology

Mice were ranked according to their ascending aortic diameter by ultrasound, and the median five per group were selected for histology. Tissue sections (10 μm) were acquired from the aortic root to the aortic arch at 100 μm intervals using a cryostat. The section corresponding to a region of maximal dilation between the sinotubular junction and the arch was analyzed. Elastin fragmentation was visualized by Verhoeff elastin staining under 20x magnification and images from three high powered fields per section were recorded for analysis. Individual data were represented as the mean of three high power fields. Fragmentation was defined as the presence of discernable breaks of continuous elastic lamina. Medial thickness was measured at the greatest thickness from inner to external elastic laminae in 3 images using NIS-Elements AR software. Measurements were verified by an independent investigator who was blinded to sample identification. More detailed descriptions of these protocols are available on protocol.io dx.doi.org/10.17504/protocols.io.be9mjh46 and dx.doi.org/10.17504/protocols.io.be9sjh6e

### AGT Western Blotting

Reducing buffer (Bio-Rad 161-0737 and Sigma M7522) and plasma (0.3 μL) from mice administered control or AGT ASO were heated to 95°C for 5 minutes. Samples were fractionated on an SDS-PAGE gel (10% wt/vol; Bio-Rad 456-8033). Proteins were transferred to a PVDF membrane via Trans-blot system (Bio-Rad 170-4256). Total proteins were detected by Ponceau S. Membranes were blocked by milk (5% wt/vol; Bio-Rad 170-6404) in TBS-T (0.1% wt/vol). Membranes were then incubated with antibodies against total AGT (0.1 μg/mL; IBL 28101) for 1 hour at room temperature then with HRP-conjugated goat-antirabbit IgG (0.2 μg/mL; Vector Pi-1000). Membranes were developed with Clarity Max ECL (Bio-Rad 1705064) on a ChemDoc MP system. Blots were quantified using Bio-Rad CFX software.

### Statistics

All animals that met pre-specified inclusion criteria, and were not excluded due to death by humane endpoint unrelated to aortic disease (fighting, infection), had cause of death adjudicated by necropsy. Statistical analyses were performed using SigmaPlot 14.0. Equal variance and normality of data determined whether non-linear, logarithmic transformation was performed and whether parametric or non-parametric tests were used. Two-way ANOVA or Student’s t-test was performed for parametric comparisons; Holm-Sidak was used for post-hoc tests. Kruskal-Wallis or Rank Sum was performed for non-parametric comparisons with Dunn’s method for post hoc tests. Data are represented as individual data points, mean ± SEM, or as box and whisker plots representing median and interquartile range where applicable.

## Results

### Progression of Aortic Dimensions was Sexually Dimorphic in Fbn1^C1041G/+^ Mice

In initial studies, the progression of aortic diameters over a 12-month interval was determined in both male and female Fbn1^+/+^ and Fbn1^C1041G/+^ mice. Because TAA in Fbn1^C1041G/+^ mice has variable pathology within the proximal thoracic aorta, several parameters were measured (**Supplemental Figure I**). This included the ascending aortic diameter, aortic root diameter, and ascending aortic length. In Fbn1^+/+^ mice, there was no statistical difference in the ascending aorta diameter, aortic root dimeter, or ascending aortic length between female and male at any interval up to 12 months of age (**Figure 1A, B, C**). At one month of age, aortic root diameters and ascending aortic lengths were increased in both male and female Fbn1^C1041G/+^ mice compared to Fbn1^+/+^ mice. However, only male Fbn1^C1041G/+^ mice exhibited statistically significant ascending aortic dilation compared to sex-matched littermates at one month of age. Despite differences at 1 month of age in female mice, the subsequent increase in diameter of ascending aorta and aortic root, and length of the ascending region were not statistically different between Fbn1^+/+^ and Fbn1^C1041G/+^ mice (**Figure 1D, E, F**). In contrast, male Fbn1^C1041G/+^ mice had augmented increases in diameters of ascending aorta and aortic root and ascending aortic length, compared to male Fbn1^+/+^ littermates over the course of 12 months. Since female Fbn1^C1041G/+^ mice had no significant differences in the progression of aortic dimensions compared to their wild type littermates, subsequent experiments used predominantly male mice.

**Figure 1:**
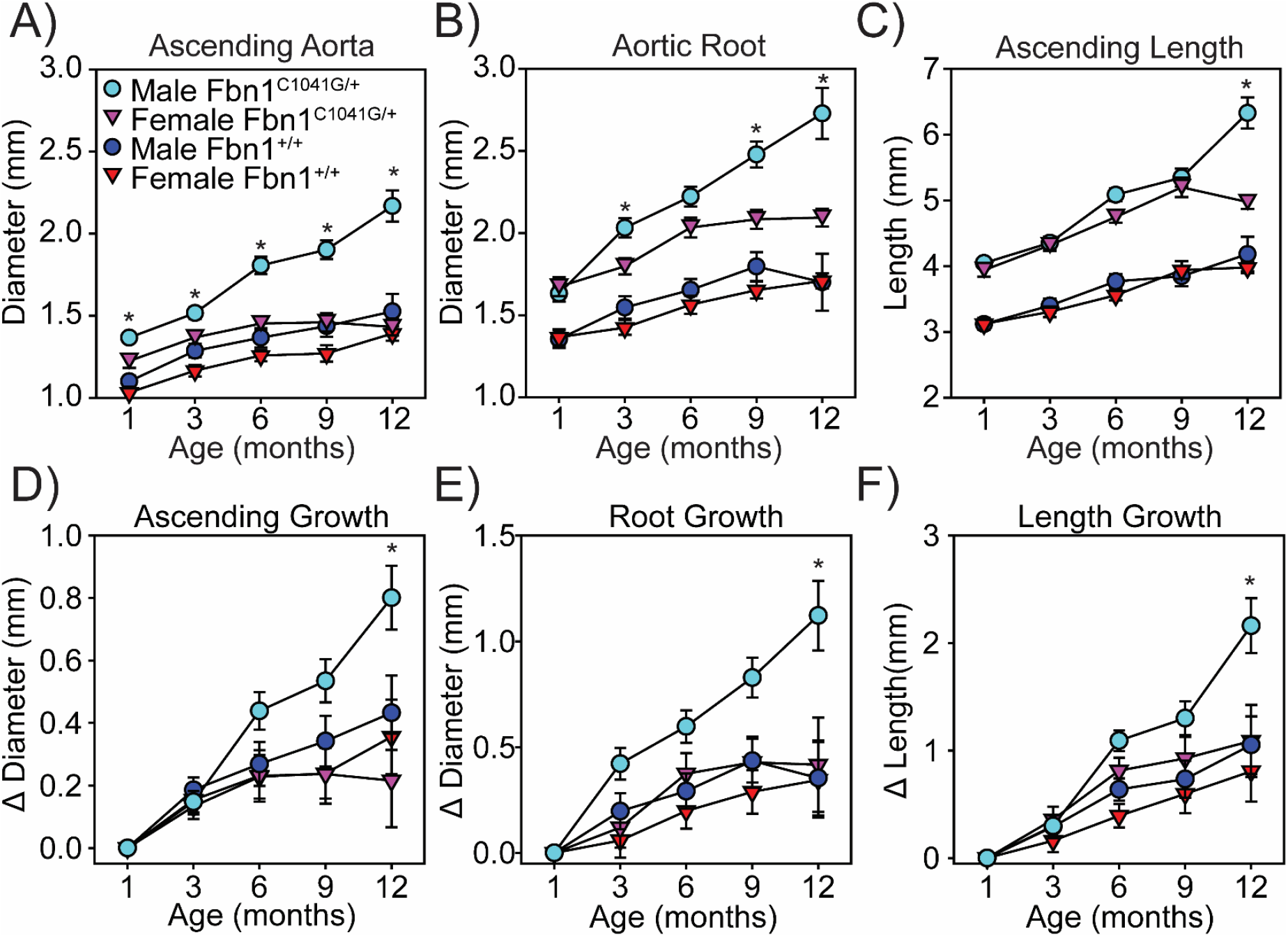
TAA in Fbn1^C1041G/+^ mice is sexually dimorphic. Sequential ultrasound measurements of the **A)** ascending aorta, **B)** aortic root, and **C)** aortic length in diastole from 1 month to 12 months of age of male and female Fbn1^+/+^ and Fbn1^C1041G/+^ mice. Data represented as change in dimensions over baseline at 1 month of age of the **D)** ascending aorta, **E)** aortic root, and **F)** aortic length. * p<0.05 of male Fbn1^C1041G/+^ versus female Fbn1^C1041G/+^ mice; n = 9-15/group.

### AT1aR Deletion Attenuated Aortic Pathology in Male Fbn1^C1041G/+^ Mice

To study the effects of AT1aR on aortic dilation in Fbn1^C1041G/+^ mice, Fbn1^C1041G/+^ mice that were either AT1aR^+/+^ or AT1aR^−/−^ were generated. Fbn1^C1041G/+^ mice were also compared against Fbn1^+/+^ mice that were also either AT1aR^+/+^ or AT1aR^−/−^. Aortic dimensions were measured using ultrasound images acquired from a right parasternal view at diastole (**Figure 2A**). Images were acquired from every mouse at the stated intervals up to 12 months of age, with no deaths of any cause occurring during the study.

**Figure 2:**
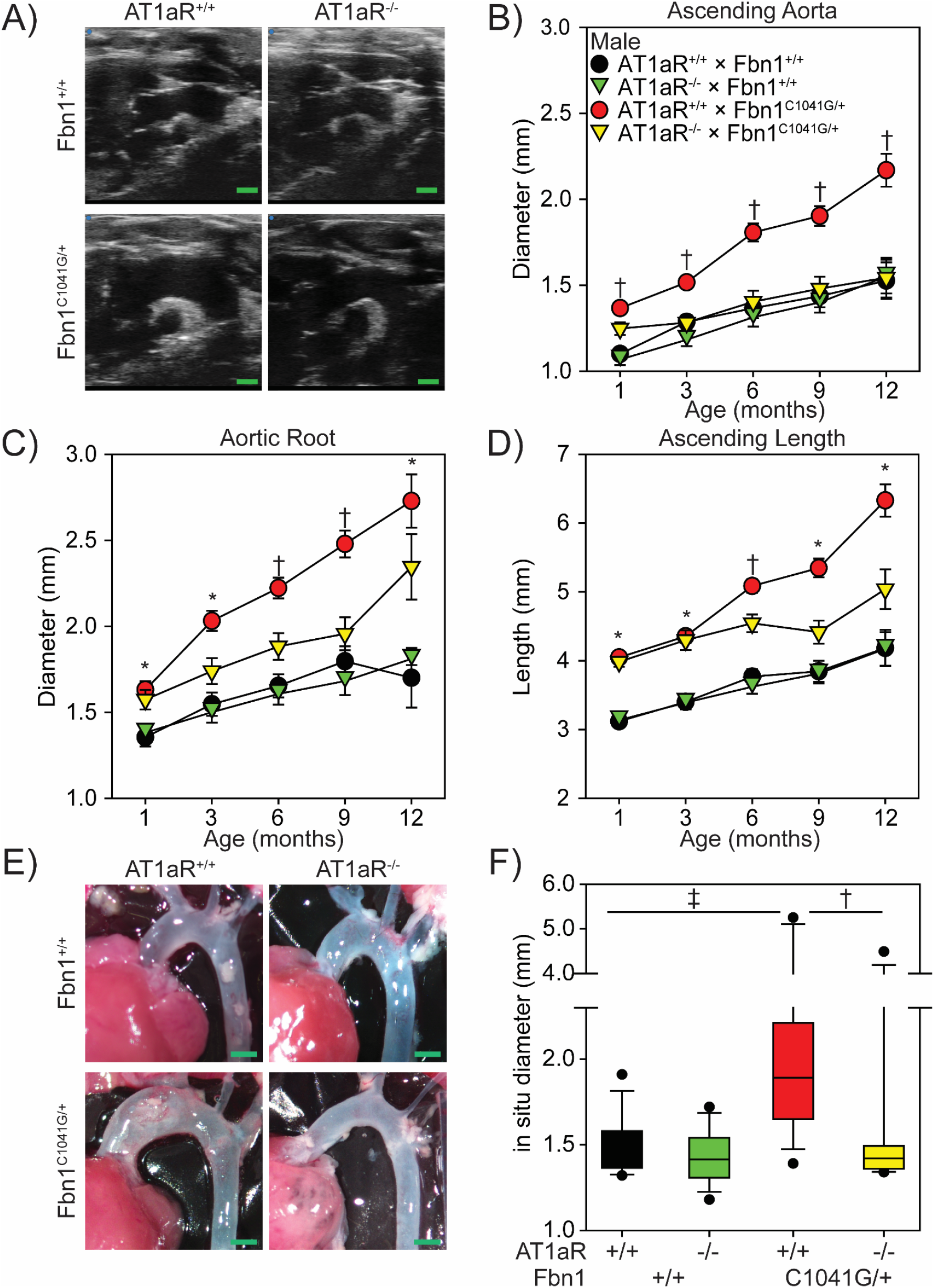
AT1aR deletion attenuated ascending aortic dilation in male Fbn1^C1041G/+^ mice. **A**) Representative ultrasound images of the thoracic aorta in male AT1aR^+/+^ x Fbn1^+/+^, AT1aR^−/−^ x Fbn1^+/+^, AT1aR^+/+^ x Fbn1^C1041G/+^, and AT1aR^−/−^ x Fbn1^C1041G/+^ mice. Green bar = 1 mm. Sequential ultrasound measurements of the **B**) ascending aorta **C**) aortic root and **D**) aortic length. * p<0.05 of AT1aR^+/+^ x Fbn1^+/+^ versus AT1aR^+/+^ x Fbn1^C1041G/+^; † p<0.05 of AT1aR^+/+^ x Fbn1^C1041G/+^ versus AT1aR^−/−^ x Fbn1^C1041G/+^; n = 11-15/group. **E**) Representative *in situ* images of the thoracic aorta. **F**) Measurement of *in situ* aortic dimensions taken at the maximal aortic diameter. † p<0.01; ‡ p<0.001; n = 10-15 / group.

Male Fbn1^+/+^ mice had modest increases in diameters of the ascending aorta (**Figure 2B**), aortic root (**Figure 2C**), and lengths of the ascending aorta (**Figure 2D**) during the course of the 12-month study. These increases were not significantly different from increases seen in Fbn1^+/+^ mice that were also AT1aR^−/−^. These findings based on the ultrasound measurements were confirmed at the 12-month interval by direct measurements on in situ aortas (**Figure 2E, F**).

At 1 month of age, male Fbn1^C1041G/+^ mice had increased diameters of ascending aorta and aortic root and lengths of ascending aorta compared to Fbn1^+/+^ mice. At this early age, deletion of AT1aR had no effect on aortic dimensions (**Supplemental Figure II**). In Fbn1^C1041G/+^ mice that were AT1aR^+/+^, there was a progressive increase in all 3 aorta dimensions acquired by ultrasound. In contrast, deletion of AT1aR markedly attenuated the progressive expansion of these dimensions to rates that were not statistically different from those in Fbn1^+/+^ mice (**Supplemental Figure III**). As with Fbn1^+/+^ mice, direct aortic measurements in situ at 12 months of age confirmed the data acquired by ultrasound. Consistent with previously publicartions,^23^ body weight and systolic blood pressure were not correlated with ascending aortic dimensions in mice (**Supplemental Figure IV**).

Female Fbn1^+/+^ and Fbn1^C1041G/+^ mice that were with either AT1aR ^+/+^ or ^−/−^ were also generated and aortic dimensions measured up to 12 months of age. As noted above, beyond the initial differences at 1 month of age, progressive changes in aortic dimensions were not different between Fbn1^+/+^ and Fbn1^C1041G/+^ female mice. The deletion of AT1aR had no effect on the age-related changes in either group (**Supplemental Figure V, VI**).

To determine if AT1aR deletion impacted the structure of the aortic media, histological characteristics were determined in aortic tissues acquired at 12 months of age. Since the most dramatic differences in changes of dimensions described above were in the ascending aorta, this region was selected for tissue characterization using our validated and reproducible method (**Supplemental Figure VII**). Ascending aortic tissues from Fbn1^+/+^ mice had elastic fibers with minimal fragmentation (**Figure 3A**). Neither the extent of fragmentation nor medial thickness were altered by the absence of AT1aR in Fbn1^+/+^ mice (**Figure 3B, C**). In contrast, Fbn1^C1041G/+^ x AT1aR^+/+^ mice had extensive fragmentation of elastic fibers and marked medial thickening. Deletion of AT1aR in these mice significantly reduced elastin fragmentation and medial thickening.

**Figure 3:**
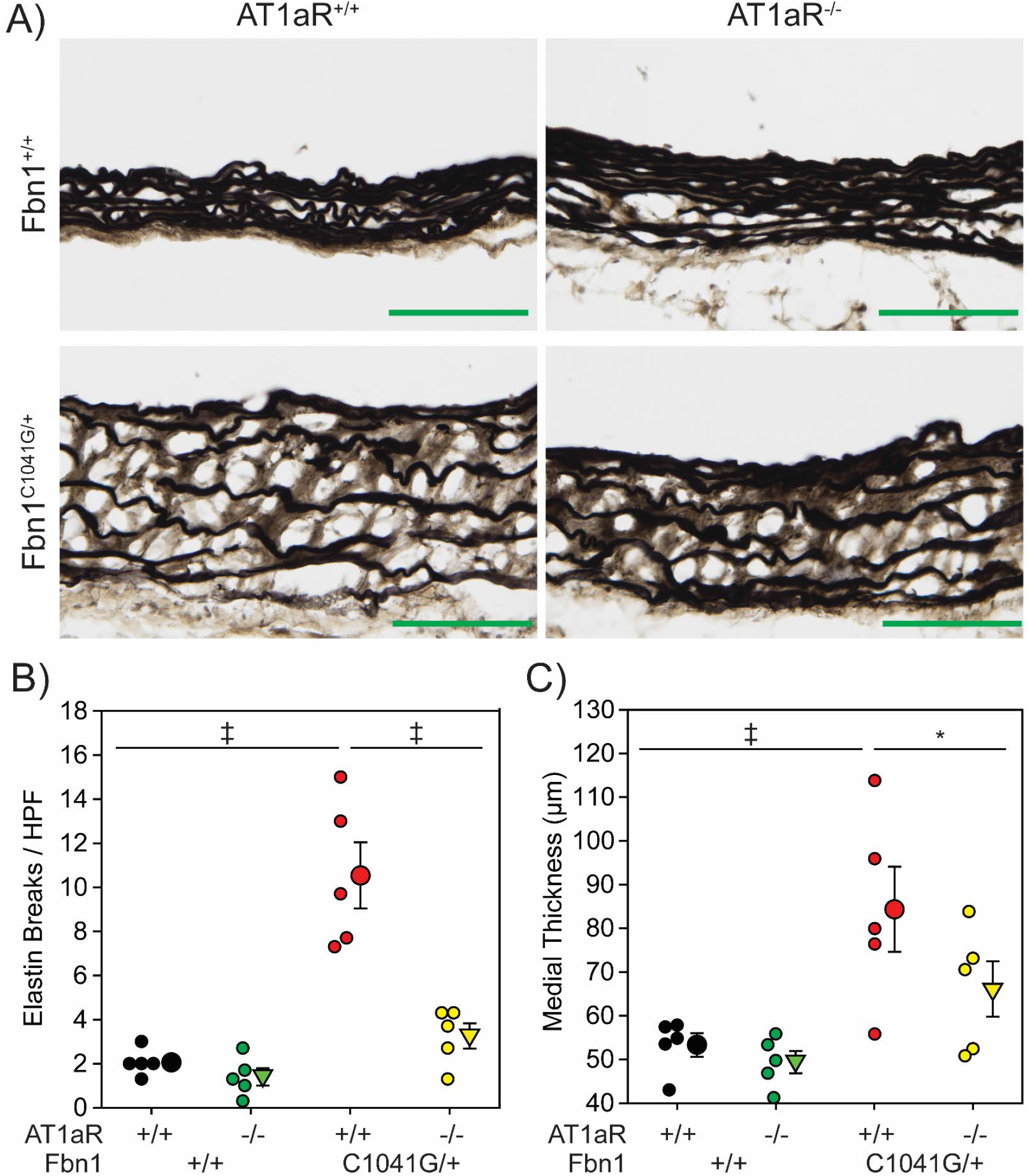
AT1aR deletion attenuated medial remodeling in male Fbn1^C1041G/+^ mice. **A**) Representative images of Verhoeff’s elastin staining in ascending aortic sections from male AT1aR^+/+^ x Fbn1^+/+^, AT1aR^−/−^ x Fbn1^+/+^, AT1aR^+/+^ x Fbn1^C1041G/+^, and AT1aR^−/−^ x Fbn1^C1041G/+^ mice. Green bar = 100 μm. **B**) Number of breaks per high powered field detected in aortic sections. **C**) Medial thickness as measured by the distance between the inner elastic lamina and external elastic lamina in aortic sections. * p<0.05, ‡ p<0.001; n = 5/group.

### Depletion of Plasma AGT Concentrations by AGT ASO Attenuated Aortic Pathology in Male Fbn1^C1041G/+^ Mice

We have demonstrated previously that administration of AGT ASO markedly reduces plasma concentration of AGT and attenuates AngII responses in mice.^24, 25^ Using ASO against the same target as previous publications, male Fbn1^C1041G/+^ mice received a loading dose (80 mg/kg) of either AGT or control ASO on day 1 and day 4 of the study. Starting on day 7, mice received a maintenance dose (40 mg/kg) every 7 days for 6 months. (**Figure 4A**). Mice tolerated the ASO well and displayed minimal toxicity after administration of loading doses (**Supplemental Figure VIII**) AGT ASO effectively depleted AGT in plasma (**Figure 4B**).

**Figure 4:**
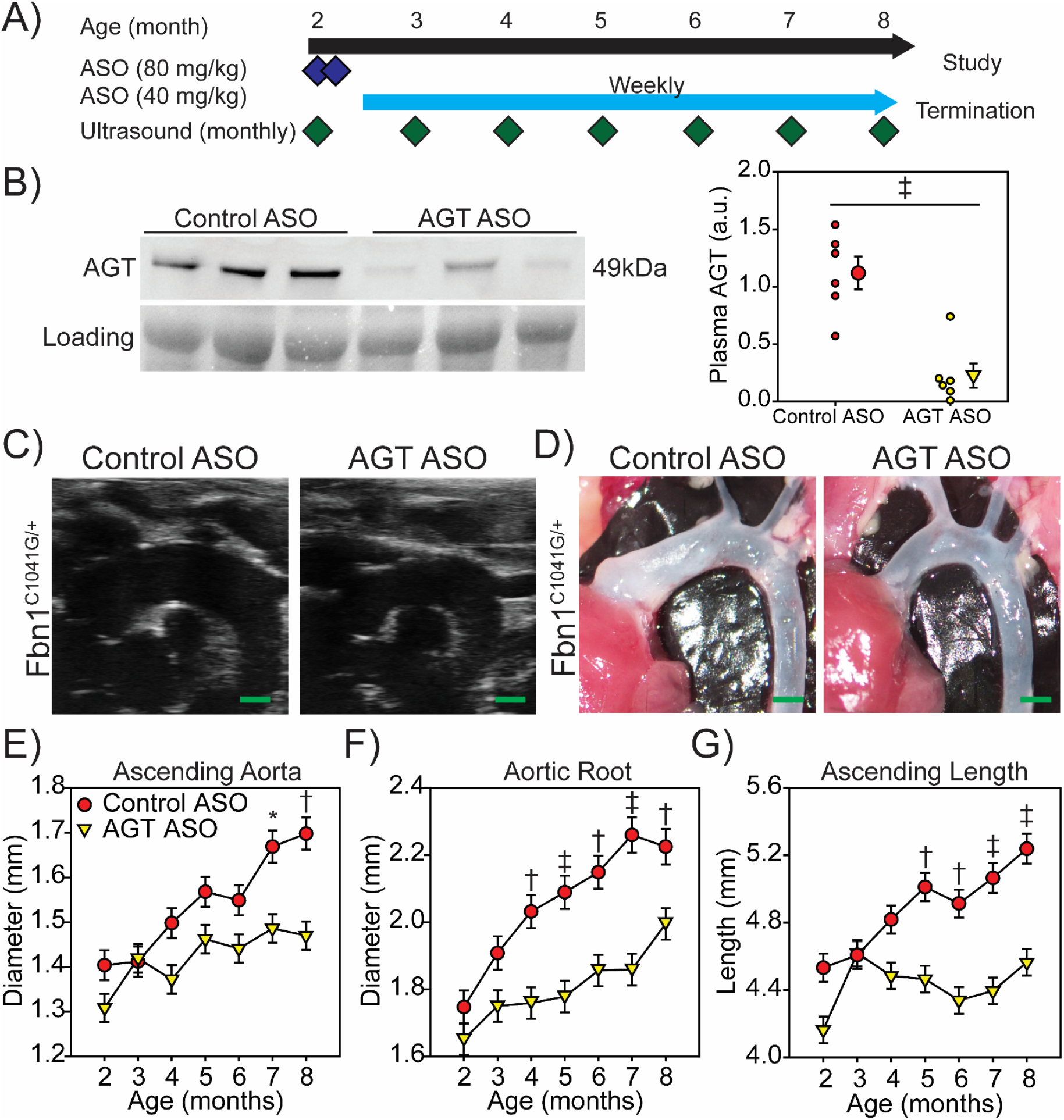
AGT ASOs depleted AGT and attenuated TAA in male Fbn1^C1041G/+^ mice. **A**) Study design and administration schedule of ASOs in male Fbn1^C1041G/+^ mice. A loading dose of control ASO or AGT ASO (80 mg/kg) was administered day 1 and 4 of study. Maintenance doses of control ASO or AGT ASO (40 mg/kg) was administered every 7 days. **B**) Western blot of plasma AGT and total plasma protein in 8 month old male Fbn1^C1041G/+^ mice administered either control ASO or AGT ASOs. Blot represents one of two experiments. ‡ p<0.001; n = 6/group. Representative **C**) ultrasound and **D**) *in situ* images of aortas from 8 month old Fbn1^C1041G/+^ mice administered either control ASO or AGT ASO. Green bar = 1 mm. Sequential ultrasound measurements of the **E**) ascending aorta, **F**) aortic root, and **G**) aortic length in diastole from 2 months to 8 months of age in male Fbn1^C1041G/+^ mice dosed with either control ASO or AGT ASO. * p<0.05, † p<0.01, ‡ p<0.001; n = 8-10/group.

Aortic dimensions were acquired starting at 2 months of age, and every month for a further 6 months using the same process described above (**Figure 4C**) with in situ aortic measurements at termination confirming the ultrasound measurement. (**Figure 4D**). AGT depletion achieved by the ASO administration led to statistically significant reductions in expansion of diameters of ascending aorta (**Figure 4E**) and aortic root (**Figure 4F**) and length of ascending aorta (**Figure 4G**) in male Fbn1^C1041G/+^ mice.

To determine whether AGT ASO impacted aortic medial structure, histology was performed on ascending aortic tissue. Consistent with our previous observation, we detected aortic medial remodeling in 8-month-old male Fbn1^C1041G/+^ mice administered control ASO (**Figure 5A**). Compared to male Fbn1^C1041G/+^ mice administered control ASO, male Fbn1^C1041G/+^ mice administered AGT ASO exhibited less elastin fragmentation and medial thickening (**Figure 5B, C**).

**Figure 5:**
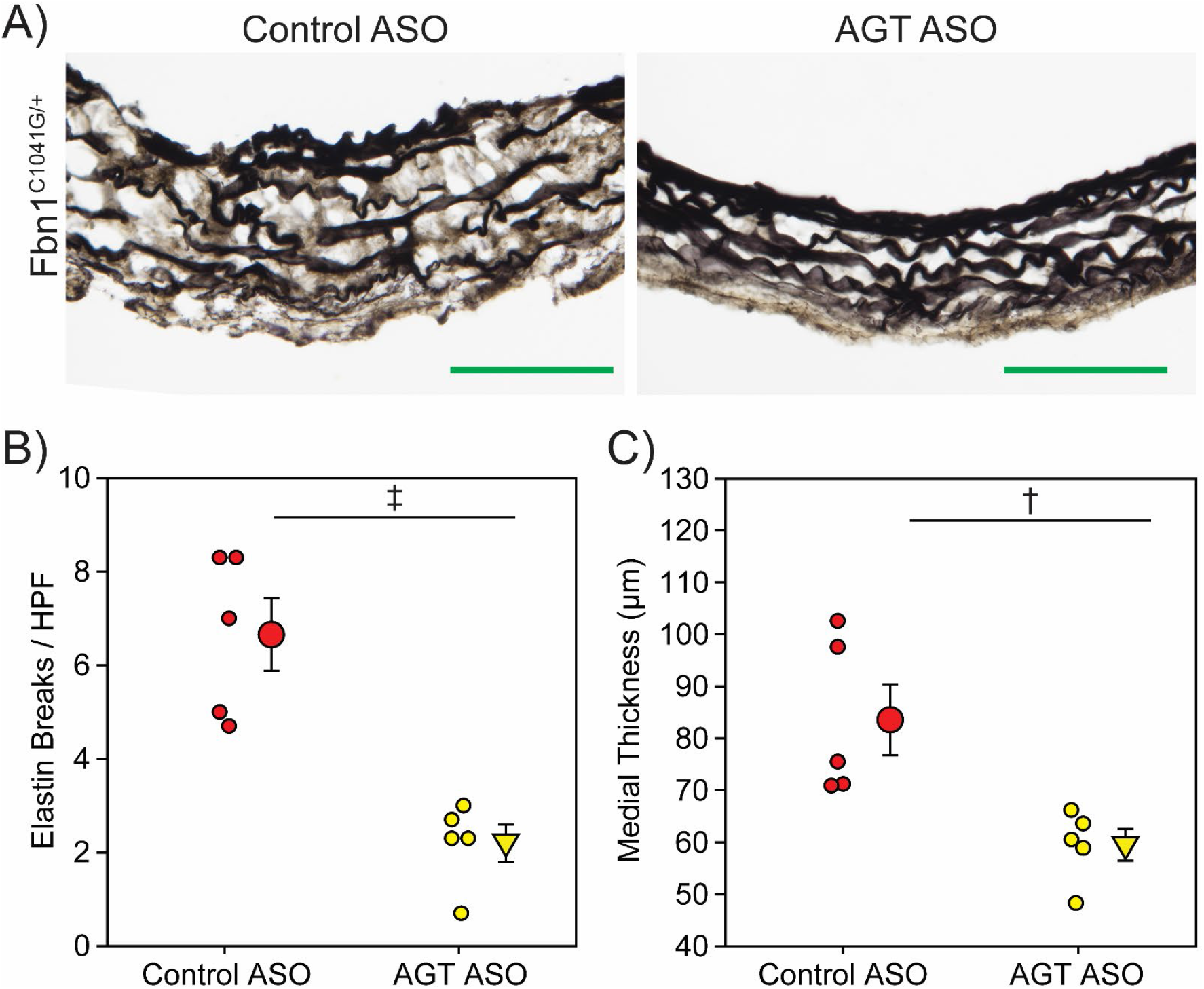
AGT ASOs attenuated medial remodeling in male Fbn1^C1041G/+^ mice. **A**) Representative images of Verhoeff’s elastin staining in aortic sections from male Fbn1^C1041G/+^ mice administered either control ASO or AGT ASO for 6 months. Green bar = 100 μm. **B**) Number of breaks per high powered field detected in aortic sections. **C**) Medial thickness as measured by the distance between the inner elastic lamina and external elastic lamina in aortic sections. † p<0.01, ‡ p<0.001; n = 5/group.

## Discussion

Using pharmacological tools to manipulate the renin angiotensin system, there have been consistent demonstrations that losartan attenuates aortic pathology in mice with fibrillin-1 manipulations.^5–8, 26–28^ However, it has been proposed that losartan may exert these beneficial actions independent of AT1 receptor antagonism.^6–8^ Additionally, it has been suggested that AngII may not be responsible for cardiovascular pathology in mice with genetically manipulated fibrillin-1.^16^ However, the present study demonstrates that both genetic deletion of AT1aR and techniques to reduce AngII availability led to reduced aortic pathology in male Fbn1^C1041G/+^ mice. These findings are consistent with ligand activation of AT1aR being the basis for aortic expansion in fibrillin-1 haplo-insufficient mice.

The sequential measurement of aortic dimensions over a protracted interval in multiple groups required development of a standardized ultrasound protocol for image acquisition. We have noted previously the variance imparted by the differences acquiring dimension at systole or diastole.^17^ Given that this excursion can be as much as 0.2 mm, lack of consistency in acquiring data could have a profound effect on data interpretation. The approach used in this study also consistently imaged the aorta from the right parasternal view.^18^ While this view is optimal for determining dimensions of the ascending aorta, it reduces precision of aortic root measurements. In the present study, there was strenuous adherence to a standardized protocol. In addition, the measurements acquired from ultrasound images were validated at termination by direct measurement of aorta in situ. This degree of measurement validation enabled acquisition of reliable data while reducing variability between sequential measurements.

Since we were not aware of any previous study that defined the effects of sex on the aortic pathology in Fbn1^C1041G/+^ mice, the initial studies used both males and females. The present study demonstrates a striking effect of sex on the aorta in these mice, with the female Fbn1^C1041G/+^ mice exhibiting minimal progression of thoracic aortic expansion compared to sex-matched Fbn1^+/+^ littermates. In mice with genetic manipulations of Fbn1, there had been only one study indicating that sexual dimorphism existed in the Fbn1^GT8/+^ mouse model of Marfan syndrome.^29^ However, sexual dimorphism of the Fbn1^GT8/+^ mouse was only defined in the context of pregnancy. In addition to revealing that female Fbn1^C1041G/+^ mice resist aortic dilation, we outlined the consequences of this sexual dimorphism on the role of AT1aR deletion. While the mechanism of this sexual dimorphism is beyond the scope of the present study, it illustrates the need for studies to report data on studies in these mice in a sex-specific manner.

Deletion of AT1aR markedly reduced progression of aortic pathology in male Fbn1^C1041G/+^ mice. AT1 receptors in mice have two isoforms, AT1aR and AT1bR, that resulted from chromosomal duplication. While there is strong sequence homology between the two isoforms, these protein have different tissue distribution and different signaling mechanisms. Absence of AT1bR has modest effects in vivo, although it is responsible for AngII induced contractions of the infrarenal mouse aorta.^30, 31^ Absence of AT1bR has no effect on AngII-induced aortopathies.^30^ AT1aR deficient mice were initially demonstrated to have lower blood pressure.^32^ However, in agreement with the present study, there has also been several publications showing no difference in blood pressure between AT1aR^+/+^ and ^−/−^ mice.^33^ While the present study was ongoing, the genetic deletion of AT1aR was reported in Fbn1 hypomorphic mice. While global deletion of AT1aR in Fbn1 hypomorphic mice had no significant effect on the survival, there was decrease aortic expansion in mice that survived to 90 days.^34^ This emphasizes the need for further study on the divergent roles of the renin angiotensin system in TAA versus aortic rupture/dissection. The early acquisition of ultrasound data in this study also illustrated that there are changes in aortic dimension in the early postnatal interval. Despite the dramatic reduction of progression of aortic dimensions in AT1aR^−/−^ mice following this postnatal interval, the absence of AT1aR failed to affect early changes. This is consistent with temporal-dependent mechanisms of the disease as have been demonstrated previously in Fbn1 hypomorphic mice.^34^

Others have noted that AT1aR deficiency had no effect on expansion of the aortic root at 3 and 6 months of age in Fbn1^C1041G/+^ mice, whereas losartan had a divergent effect and was able to decrease aortic root expansion in these mice.^6^ The beneficial effects of losartan were attributed to preservation of endothelial function in an AT1aR independent manner through an alternative VEGFR2/eNOS pathway. The basis for the disparity relative to the present study are not clear. Comparisons are hampered by the paucity of data on the protocol for ultrasound acquisition and on the sex of the mice in each group. Other studies suggested that losartan’s protective effect may be due to tumor growth factor β inhibition or AngII receptor type 2 hyperstimulation.^5, 8^ However, our data indicated that blockade of the AT1aR is sufficient to attenuate Marfan syndrome associated TAA. While the pleotropic effects of losartan may contribute to attenuating thoracic aortopathies, the present study is consistent with the postulate that its benefit is due to inhibition of AT1aR activation.

We used an ASO to decrease the synthesis of the unique precursor of all angiotensin peptides to determine whether AT1aR stimulation in aortopathies required AngII as a ligand. This approach is advantageous over the more common mode of reducing AngII production through inhibiting angiotensin-converting enzyme, which regulates other pathways including the kinin-kallikrein system. Additionally, the protracted half-life of ASO leads to persistent inhibition of AGT synthesis and profound reductions in plasma AGT concentrations. Use of this pharmacologic modality also avoids adverse consequences of genetic deletion of the renin angiotensin system components. Previous genetic approaches have included the use of mice with global deficiencies of AGT. However, these mice have several major developmental abnormalities include poor growth and cardiomyopathy.^35^ Inhibition of AGT synthesis by an ASO reduces plasma concentrations by approximately 90% in the postnatal phase with no observable toxicity as demonstrated in the present study and other reports.^24, 36^ Therefore, the use of ASO to deplete AGT demonstrated the need for the presence of angiotensin ligands to augment aortic pathology in Fbn1^C1041G/+^ mice.

In humans, randomized control trials of angiotensin receptor blockers have yielded mixed results in Marfan syndrome associated TAA, in contrast to the consistent results that have been generated using mouse models of the disease.^5–8, 26–28^ Most of the mouse and human studies have been performed using losartan, which is characterized by a relatively short half-life and surmountable antagonism. The deficiencies of this drug were likely to have been ameliorated in mouse studies by consistent delivery, via osmotic pumps and diet, leading to a persistent inhibition. AT1 receptor antagonists with enhanced pharmacological profiles, such as irbesartan and candesartan, would be preferable to test the role of AT1 receptor inhibition in humans. Indeed, it has been demonstrated recently that irbesartan significantly attenuated aortic root expansion in individuals with Marfan syndrome.^37^ Conversely, ASO affords chronic and persistent inhibition of AGT synthesis to effect long-term depletion of angiotensin ligands. These durable effects of ASO enables inhibition of AGT synthesis to be tested as a possible approach to reduce TAA in Marfan syndrome.

Our study provided strong evidence that both AT1aR deletion and AGT depletion resulted in significant attenuation of ascending aortic dilation and lengthening. These data are consistent with AngII signaling through AT1aR being necessary for TAA progression in male Fbn1^C1041G/+^ mice and that profound and persistent depletion of either component is sufficient to attenuate TAA. This study enables future studies to focus on cell types(s) expressing AT1aR that are stimulated to promote the disease. These studies would give great insight into the role of AT1aR on key spatial and temporal events during TAA development but would require generation of cell-specific and lineage-traced AT1aR knockouts in a Marfan mouse model. In addition, this study outlines the need to use pharmacologic agents that have durable inhibition of the renin angiotensin system. Durable inhibition would encompass use of angiotensin receptor blockers with long half-lives and unsurmountable modes of inhibition as well as ASO based approaches. Collectively, these data indicate that renin angiotensin system blockade holds promise in treating Marfan syndrome associated TAA when durable inhibition is achieved.

## Non-standard abbreviations

TAA: Thoracic Aortic Aneurysm
AT1aR: Angiotensin II Receptor Type 1a
Fbn1: Fibrillin-1
AGT: Angiotensinogen
AngII: Angiotensin II
ASO: Antisense Oligonucleotide

## Acknowledgments

We thank the University of Kentucky – Baylor College of Medicine Aortic Research Center investigators for their input in this manuscript.

## Sources of Funding

The authors’ research work is supported by the American Heart Association SFRN in Vascular Disease (18SFRN33960163) and the NIH NHLBI under award numbers R01HL133723. JC and MS have been supported by NCATS UL1TR001998 and JC has been supported by NHLBI F30143943. HS is supported by an AHA postdoctoral fellowship (18POST33990468). The content in this manuscript is solely the responsibility of the authors and does not necessarily represent the official views of the American Heart Association or the National Institutes of Health.

## Disclosures

MS, AD, HL, and JC have submitted a patent application for use of antisense oligonucleotides targeted against angiotensinogen in thoracic aneurysmal disease. AM is an employee of Ionis Pharmaceuticals, who provided the Control ASO and AGT ASO. Ionis Pharmaceuticals did not provide funding for this study.

## SUPPLEMENTAL DATA

**Supplemental Figure I:**
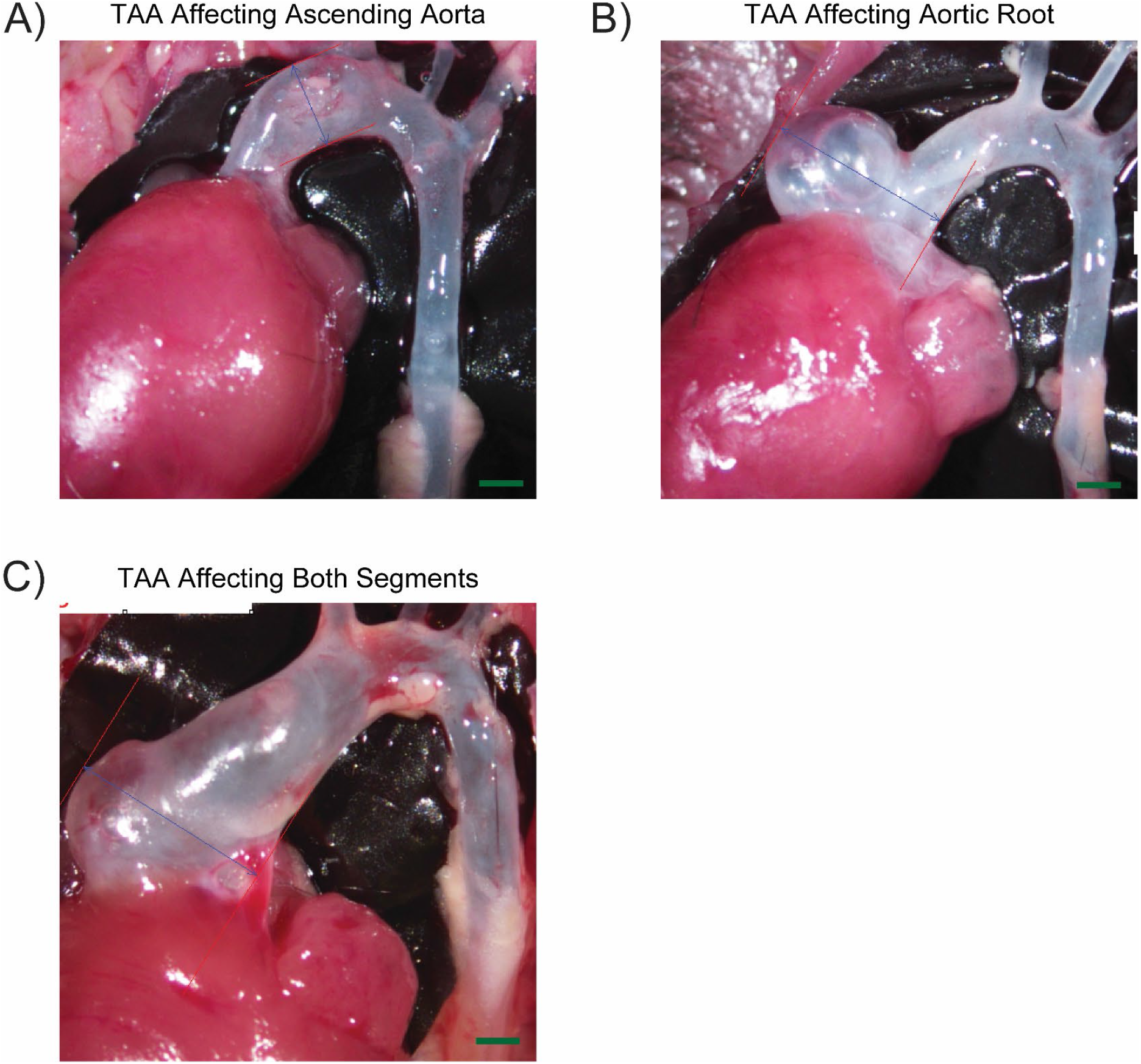
Regional heterogeneity of TAA in Fbn1^C1041G/+^ mice. TAAs in one year old Fbn1^C1041G/+^ mice have a variable phenotype of pathology location. Examples include aneurysmal presence in; **A**) ascending aorta only, **B**) aortic root only, or **C**) both segments. Green bar = 1 mm.

**Supplemental Figure II:**
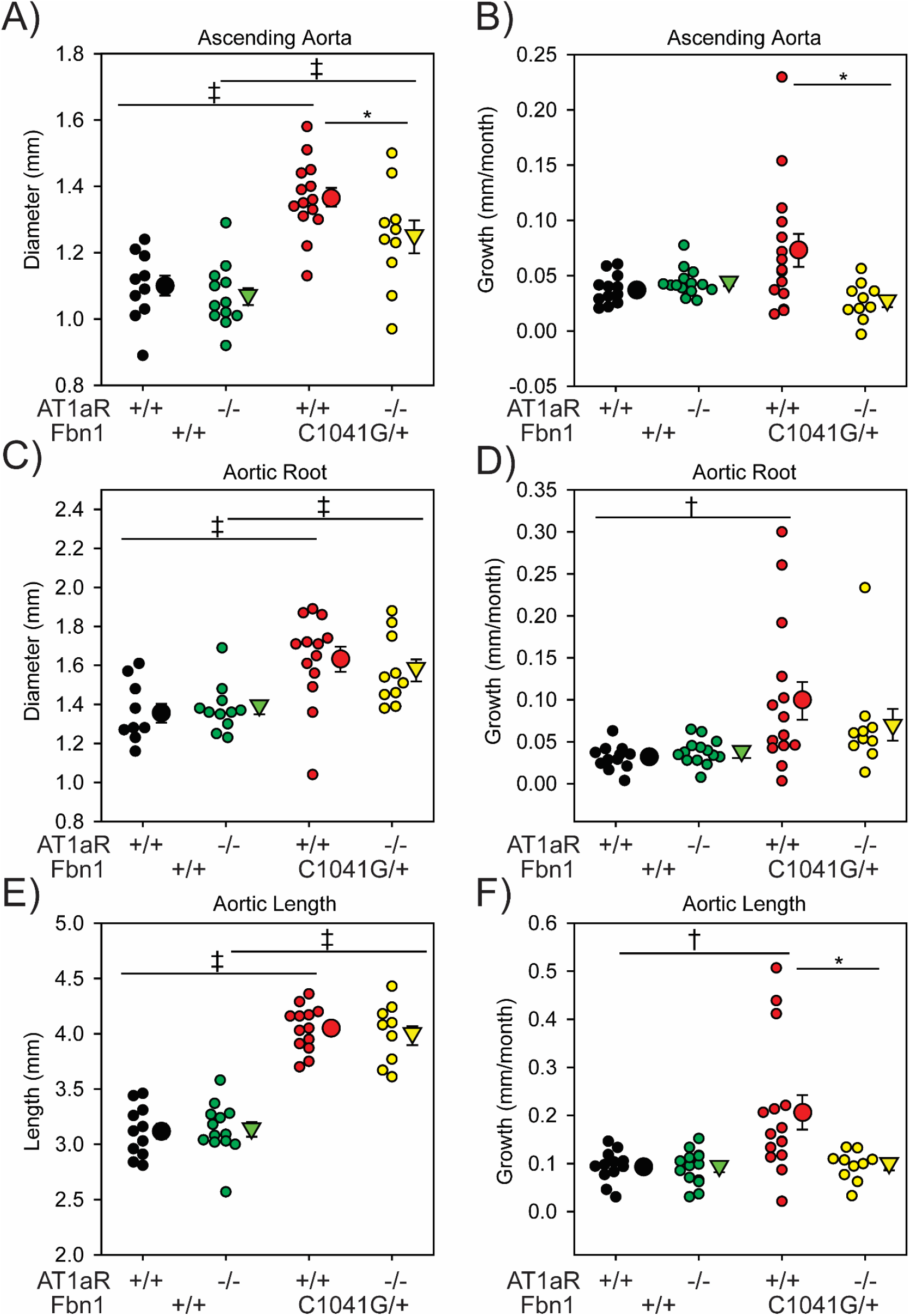
Aortic dimensions at 1 month of age and aortic growth in male mice. Ultrasound measurements of the **A**) ascending aorta, **C**) aortic root, and **E**) aortic length in diastole at 1 month of age from male AT1aR^+/+^ x Fbn1^+/+^, AT1aR^−/−^ x Fbn1^+/+^, AT1aR^+/+^ x Fbn1^C1041G/+^, and AT1aR^−/−^ x Fbn1^C1041G/+^ mice. Mean monthly ascending **B**) ascending aorta growth, **D**) aortic root growth, and **F**) aortic length growth from 1 month to 12 months in male AT1aR^+/+^ x Fbn1^+/+^, AT1aR^−/−^ x Fbn1^+/+^, AT1aR^+/+^ x Fbn1^C1041G/+^, and AT1aR^−/−^ x Fbn1^C1041G/+^ mice. * p<0.05, † p<0.01, ‡ p<0.001; n = 11-15/group.

**Supplemental Figure III:**
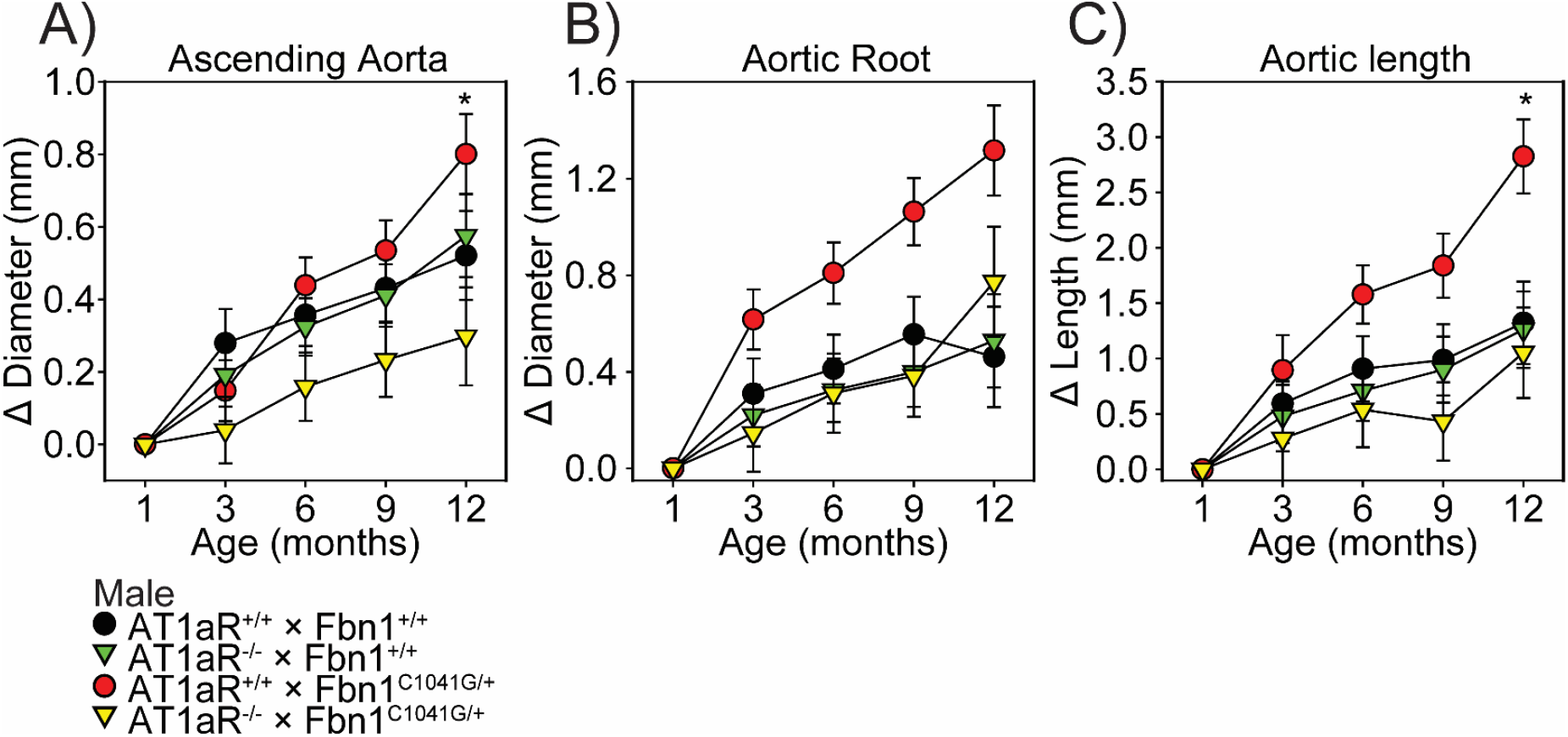
Growth from 1 month of age in male AT1aR deleted, Fbn1^C1041G/+^ mice. Data are represented as change in dimensions over the measurement at 1 month of age of the **A**) ascending aorta, **B**) aortic root, and **C**) aortic length. * p<0.05 of AT1aR^+/+^ x Fbn1^C1041G/+^ versus AT1aR^−/−^ x Fbn1^C1041G/+^; n = 11-11/group.

**Supplemental Figure IV:**
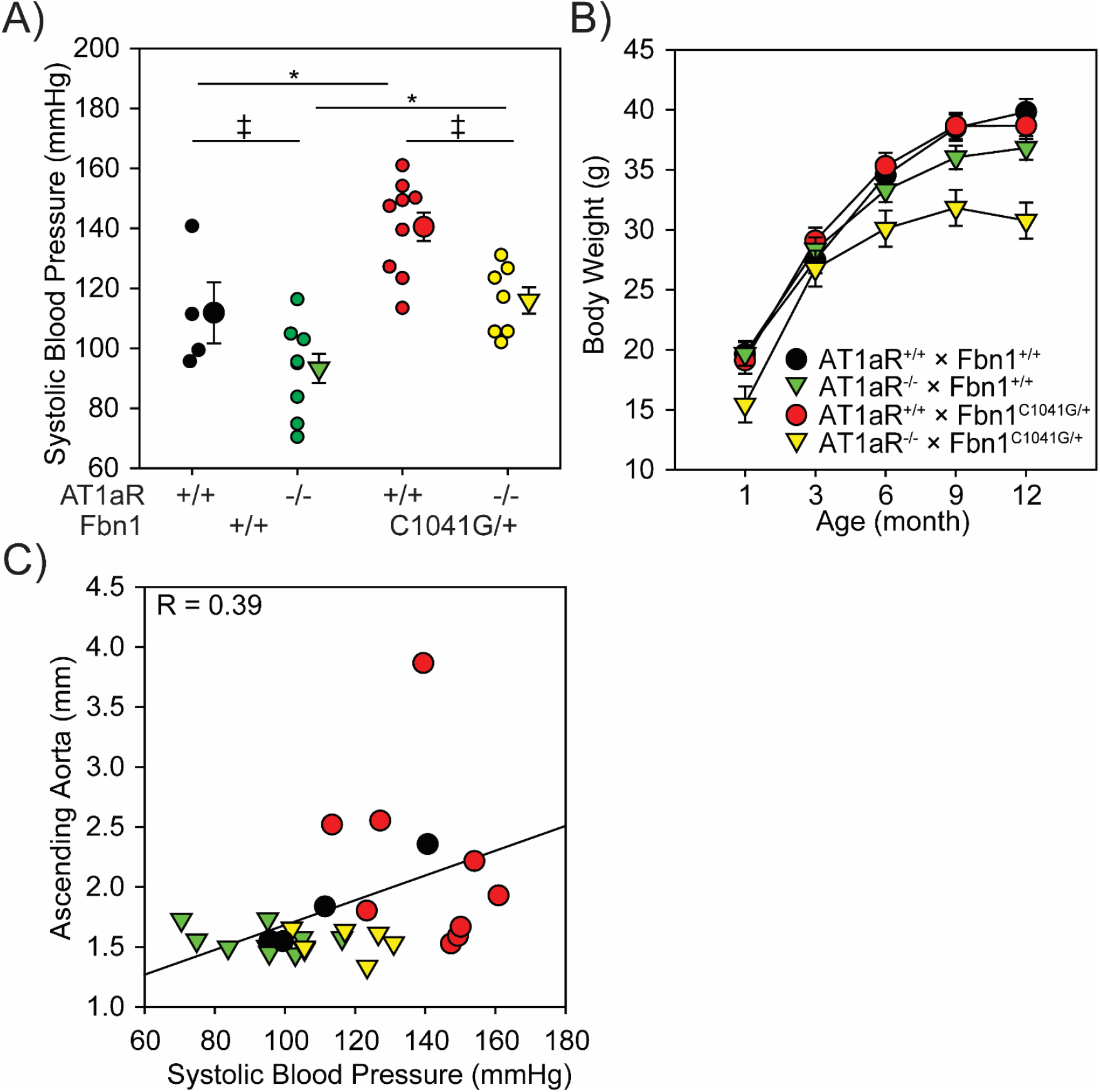
Confounding factors did not contribute to TAA phenotype in male Fbn1^C1041G/+^ mice. **A**) Systolic blood pressure measured by a tail cuff based technique in 12 month old male mice. * p<0.05, † p<0.01, ‡ p<0.001; n = 5-10/group. **B**) Sequential body weight of male mice. **C**) Correlation between systolic blood pressure and aortic diameters at 12 months of age between male mice. n = 5-10/group. Black = AT1aR^+/+^ x Fbn1^+/+^, green = AT1aR^−/−^ x Fbn1^+/+^, red = AT1aR^+/+^ x Fbn1^C1041G/+^, and yellow = AT1aR^−/−^ x Fbn1^C1041G/+^.

**Supplemental Figure V:**
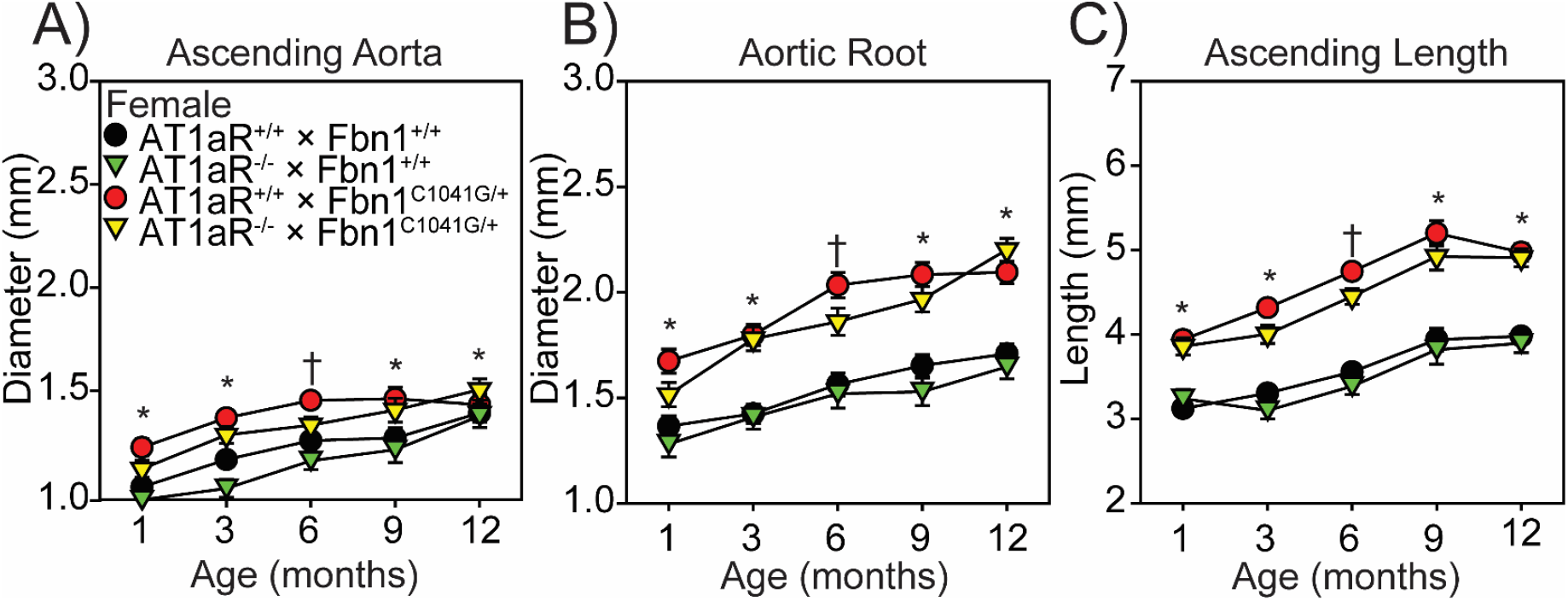
AT1aR deletion had no effect on aortic measurements in female Fbn1^C1041G/+^ mice. Sequential ultrasound measurements of the; **A**) ascending aorta, **B**) aortic root, and **C**) aortic length in diastole from 1 month to 12 months of age of female AT1aR^+/+^ x Fbn1^+/+^, AT1aR^−/−^ x Fbn1^+/+^, AT1aR^+/+^ x Fbn1^C1041G/+^, and AT1aR^−/−^ x Fbn1^C1041G/+^ mice. * p<0.05 of AT1aR^+/+^ x Fbn1^+/+^ versus AT1aR^+/+^ x Fbn1^C1041G/+^; † p<0.05 of AT1aR^+/+^ x Fbn1^C1041G/+^ versus AT1aR^−/−^ x Fbn1^C1041G/+^; n = 7-11/group.

**Supplemental Figure VI:**
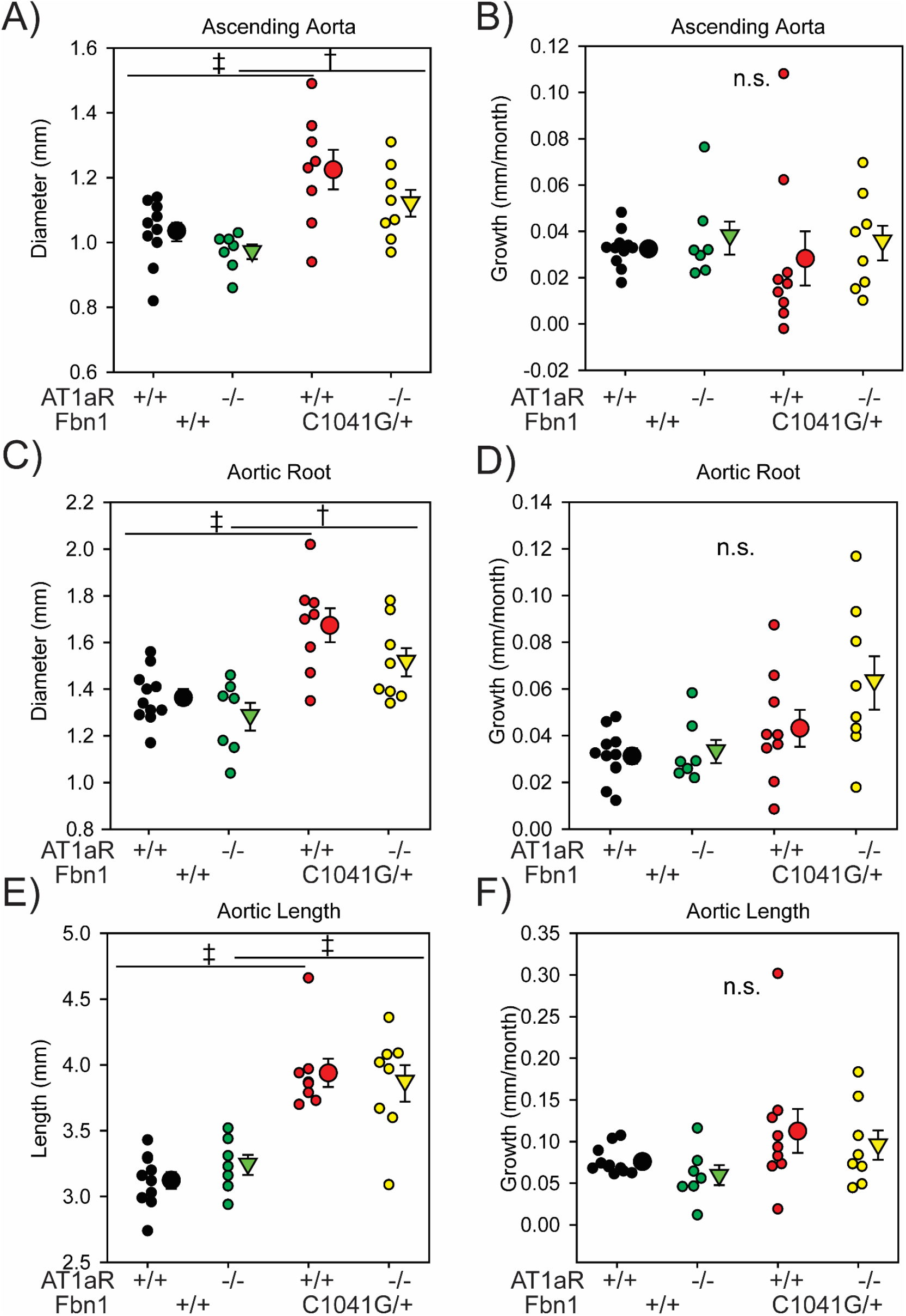
Aortic dimensions at 1 month of age and aortic growth in female mice. Ultrasound measurements of the; **A**) ascending aorta, **C**) aortic root, and **E**) aortic length in diastole at 1 month of age from female AT1aR^+/+^ x Fbn1^+/+^, AT1aR^−/−^ x Fbn1^+/+^, AT1aR^+/+^ x Fbn1^C1041G/+^, and AT1aR^−/−^ x Fbn1^C1041G/+^ mice. Mean monthly ascending **B**) ascending aorta growth, **D**) aortic root growth, and **F**) aortic length growth from 1 month to 12 months in female AT1aR^+/+^ x Fbn1^+/+^, AT1aR^−/−^ x Fbn1^+/+^, AT1aR^+/+^ x Fbn1^C1041G/+^, and AT1aR^−/−^ x Fbn1^C1041G/+^ mice. * p<0.05, † p<0.01, ‡ p<0.001; n = 7-11/group.

**Supplemental Figure VII:**
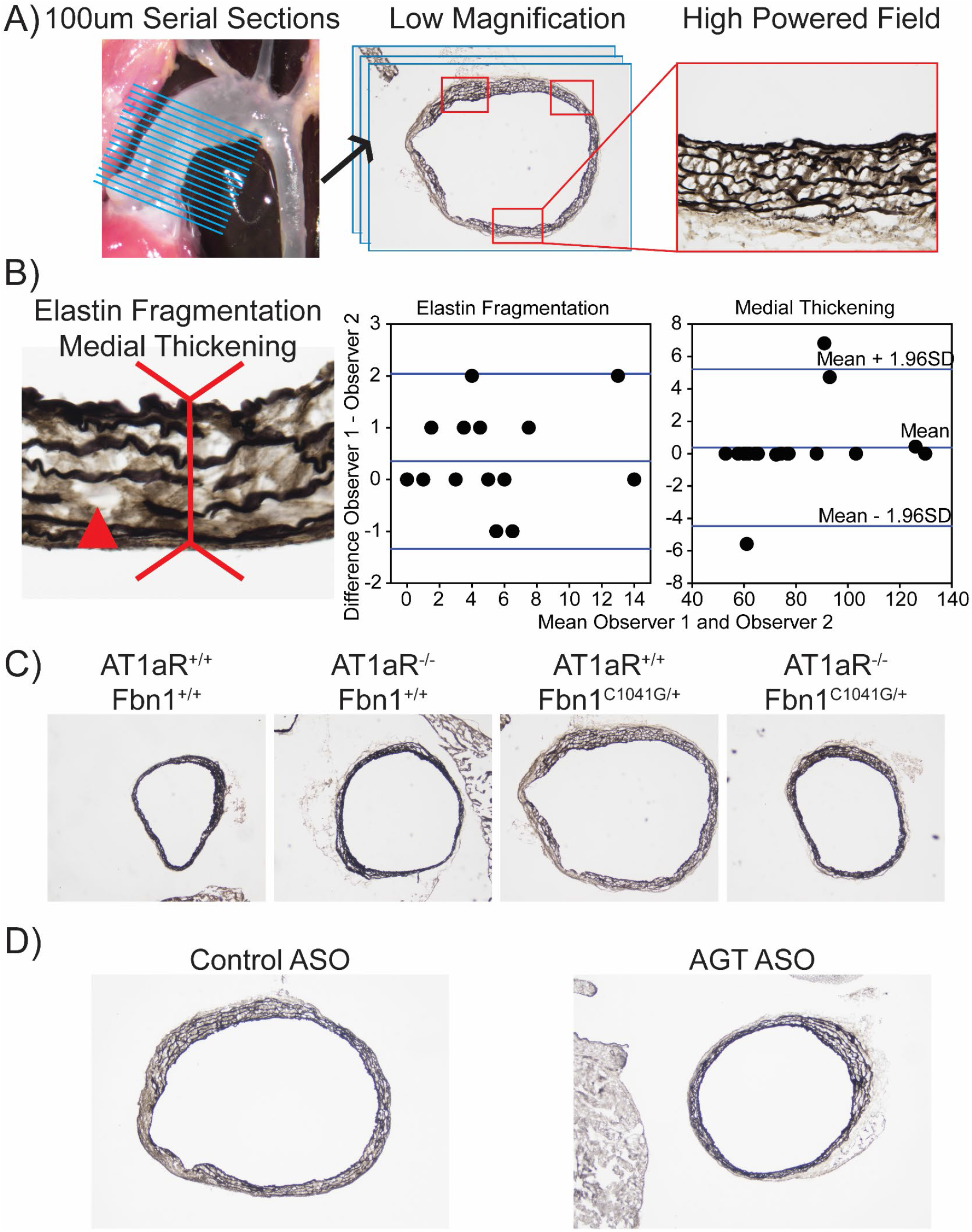
Generation of ascending aortic sections to measure elastin fragmentation and medial thickening. **A**) Generation of serial sections of ascending aortas used for histology. (blue lines) Three high powered fields / section were imaged and quantified per biological replicate. (red boxes) **B**) Quantification of elastin fragmentation (triangle) and medial thickening (inverted double arrow) by two independent observers demonstrated good agreement in both measures via Bland-Altman analysis. **C**) Low magnification images of aortic sections stained with Verhoeff elastin stain in 12 month old male AT1aR^+/+^ x Fbn1^+/+^, AT1aR^−/−^ x Fbn1^+/+^, AT1aR^+/+^ x Fbn1^C1041G/+^, and AT1aR^−/−^ x Fbn1^C1041G/+^ mice. **D**) Low magnification images of aortic sections stained with Verhoeff elastin stain in 8 month old male Fbn1^C1041G/+^ mice after 6 months of control ASO or AGT ASO.

**Supplemental Figure VIII:**
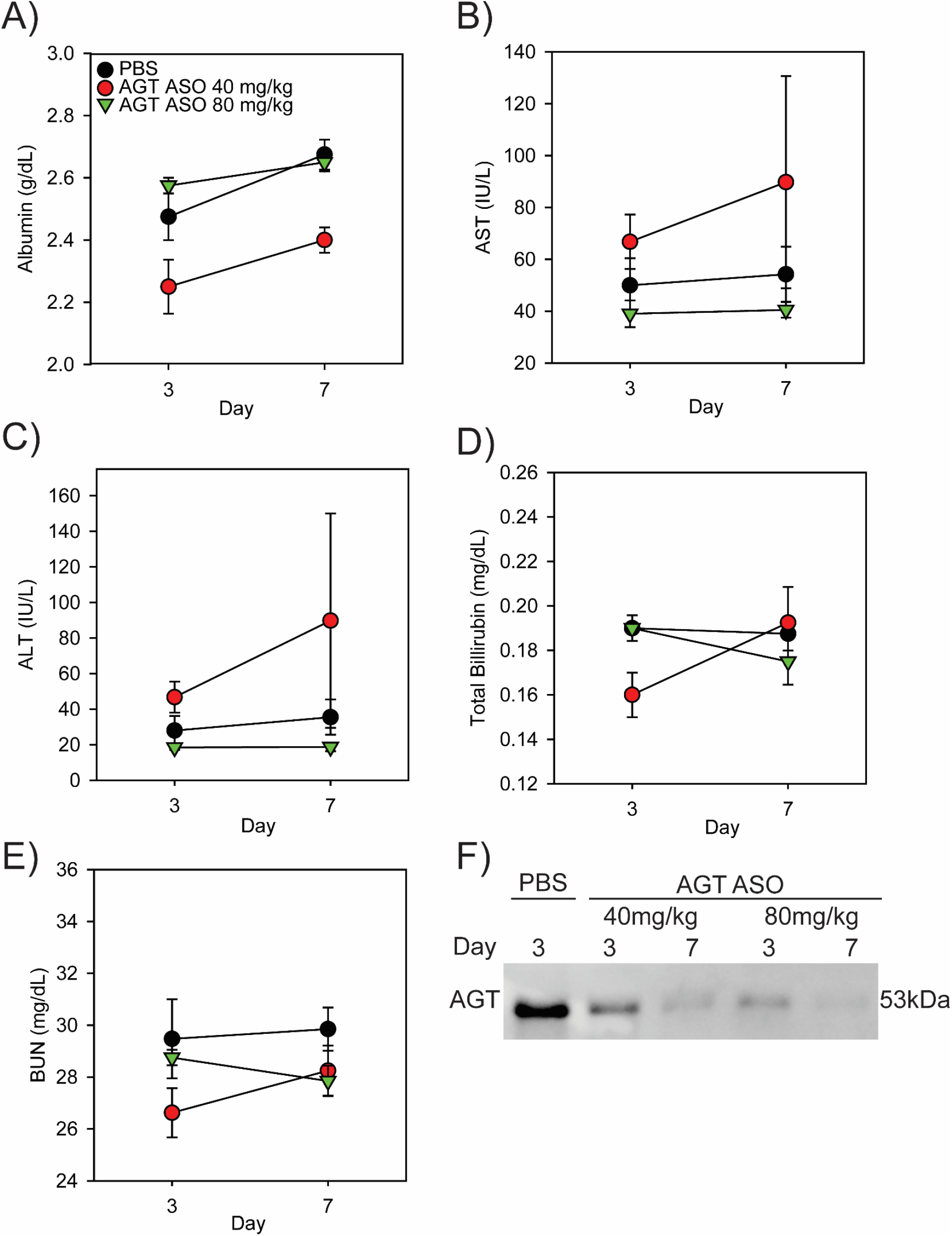
AGT ASOs have low toxicity and effectively reduce circulating AGT. Plasma concentrations of; **A**) albumin, **B**) alanine transaminase (AST), **C**) aspartate aminotransferase (ALT), **D**) total bilirubin **E**) blood urea nitrogen (BUN) in mice administered either PBS, AGT ASO (40 mg/kg), or AGT ASO (80 mg/kg) at days 1 and 4. Plasma was taken at days 3 and 7. n = 4/group. **F**) Plasma AGT concentrations at days 3 and 7 detected by Western blotting after AGT ASO was administered at days 1 and 4.

## Major Resources Table

In order to allow validation and replication of experiments, all essential research materials listed in the Methods should be included in the Major Resources Table below. Authors are encouraged to use public repositories for protocols, data, code, and other materials and provide persistent identifiers and/or links to repositories when available. Authors may add or delete rows as needed.

Animals (in vivo studies)
| Species | Vendor or Source | Background Strain | Sex | Persistent ID / URL |
| --- | --- | --- | --- | --- |
| Mice | The Jackson Laboratory | C57BL/6J | M / F | See below |

Genetically Modified Animals
|  | Species | Vendor or Source | Background Strain | Other Information | Persistent ID / URL |
| --- | --- | --- | --- | --- | --- |
| Parent – Male | Mice | The Jackson Laboratory | C57BL/6J | (B6.129-Fbn1tm1Hcd/J) FBN1<sup>C1041G/+</sup> | # 012885 |
| Parent – Female | Mice | The Jackson Laboratory | C57BL/6J | (B6.129P2-Agtr1atm1Unc/J) AT1aR<sup>-/-</sup> | # 002682 |

Antibodies
| Target antigen | Vendor or Source | Catalog # | Working concentration | Lot # (preferred but not required) | Persistent ID / URL |
| --- | --- | --- | --- | --- | --- |
| Total AGT | IBL | 28101 | 0.1 µg/mL | 1L-325 | AB 2034701 |
| HRP-conjugated goat-antirabbit IgG | Vector | Pi-1000 | 0.2 µg/mL | X0126 | AB 2336198 |

DNA/cDNA Clones
| Clone Name | Sequence | Source / Repository | Persistent ID / URL |
| --- | --- | --- | --- |
| N/A |  |  |  |

Cultured Cells
| Name | Vendor or Source | Sex (F, M, or unknown) | Persistent ID / URL |
| --- | --- | --- | --- |
| N/A |  |  |  |

Data & Code Availability
| Description | Source / Repository | Persistent ID / URL |
| --- | --- | --- |
| N/A |  |  |

Other
| Description | Source / Repository | Persistent ID / URL |
| --- | --- | --- |
| N/A |  |  |

